# Lactate Increases Stemness of CD8^+^ T Cells to Augment Anti-Tumor Immunity

**DOI:** 10.1101/2021.10.18.464724

**Authors:** Qiang Feng, Zhida Liu, Xuexin Yu, Tongyi Huang, Jiahui Chen, Jian Wang, Jonathan Wilhelm, Suxin Li, Jiwon Song, Wei Li, Zhichen Sun, Baran Sumer, Bo Li, Yang-Xin Fu, Jinming Gao

## Abstract

Nutrients and metabolites play important roles in immune functions. Recent studies show lactate instead of glucose can serve as a primary carbon fuel source for most tissues. The role of lactate in tumor immunity is not well understood with immune suppressive functions reported for lactic acid, the conjugate acid form of lactate. In this study, we report lactate increases the stemness of CD8^+^ T cells and augments anti-tumor immunity. Subcutaneous administration of lactate but not glucose shows CD8^+^ T cell-dependent tumor growth inhibition. Single cell transcriptomics analysis revealed lactate treatment increased a subpopulation of stem-like TCF-1-expressing CD8^+^ T cells, which is further validated by *ex vivo* culture of CD8^+^ T cells from mouse splenocytes and human peripheral blood mononuclear cells. The inhibition of histone deacetylase activity by lactate increased acetylation in the histone H3K27 site at the *Tcf7* super enhancer locus and increased the gene expression of *Tcf7*. Adoptive transfer of CD8^+^ T cells pretreated with lactate *in vitro* showed potent tumor growth inhibition *in vivo*. Our results elucidate the immune protective role of lactate in anti-tumor immunity without the masking effect of acid. These results may have broad implications for T cell therapy and the understanding of lactate in immune metabolism.

## Introduction

Extensive efforts have recently been dedicated to the investigation of metabolites and metabolic processes in the regulation of immune functions ^1-3^. Various metabolites (e.g., glucose, fatty acid, amino acid) coordinate with immune signaling pathways to control immune cell function and differentiation ^4-8^. Metabolic reprogramming is emerging as a new therapeutic principle to augment immune functions and immunotherapy outcomes ^9, 10^.

Lactate has historically been known as a metabolic waste product from fermentation of carbohydrates or anaerobic glycolysis in skeletal muscles during exercise ^11, 12^. Over a century ago, Otto Warburg noted cancer cells rapidly produce lactic acid even in the presence of oxygen, a process known as aerobic glycolysis ^13, 14^. Recent studies in human lung cancer patients show lactate can be used as an energy source by the cancer cells ^15, 16^. Further reports demonstrate lactate can overtake glucose as a primary carbon fuel source for a majority of tissues including immune organs ^17, 18^. The effect of lactate on CD8^+^ T cell immune functions is not well understood with immune suppressive functions reported for lactic acid ^19-21^. In this study, we investigated the effect of lactate, apart from its acidic counterpart, on immune functions and uncovered an immune protective role through the boosting of stem-like CD8^+^ T cells in cancer treatment.

## Results

### Lactate promotes antitumor immunity through CD8^+^ T cells in multiple tumor models

To elucidate the effect of lactate on anti-tumor immune response, we treated tumor bearing mice with subcutaneous (s.c.) administration of sodium lactate solution (pH 7.4, 1.7 g/kg, **Fig. 1a**). Glucose solution (pH 7.4, 5 g/kg) was used as a control. In the MC38 colon cancer model, lactate treatment suppressed tumor growth significantly, while in contrast, glucose had minimal effect (**Fig. 1b**). The immune cell dependence was tested on three major immune cell populations (CD8^+^ T cells, CD4^+^ T cells and macrophages) in tumor microenvironment. Lactate treatment shows no effect on the tumor growth in *Rag1*^-/-^ mice, indicating that T cells are necessary for the lactate-induced tumor growth inhibition (**Fig. 1c**). Depletion of CD8^+^ T cells abolished the antitumor effect of lactate (**Fig. 1d**). In contrast, the anti-tumor efficacy of lactate was not affected by the blockings of CD4^+^ T cells or macrophages (**Fig. 1e,f**). These results clarify that lactate promotes anti-tumor immunity through CD8^+^ T cells.

**Fig. 1.**
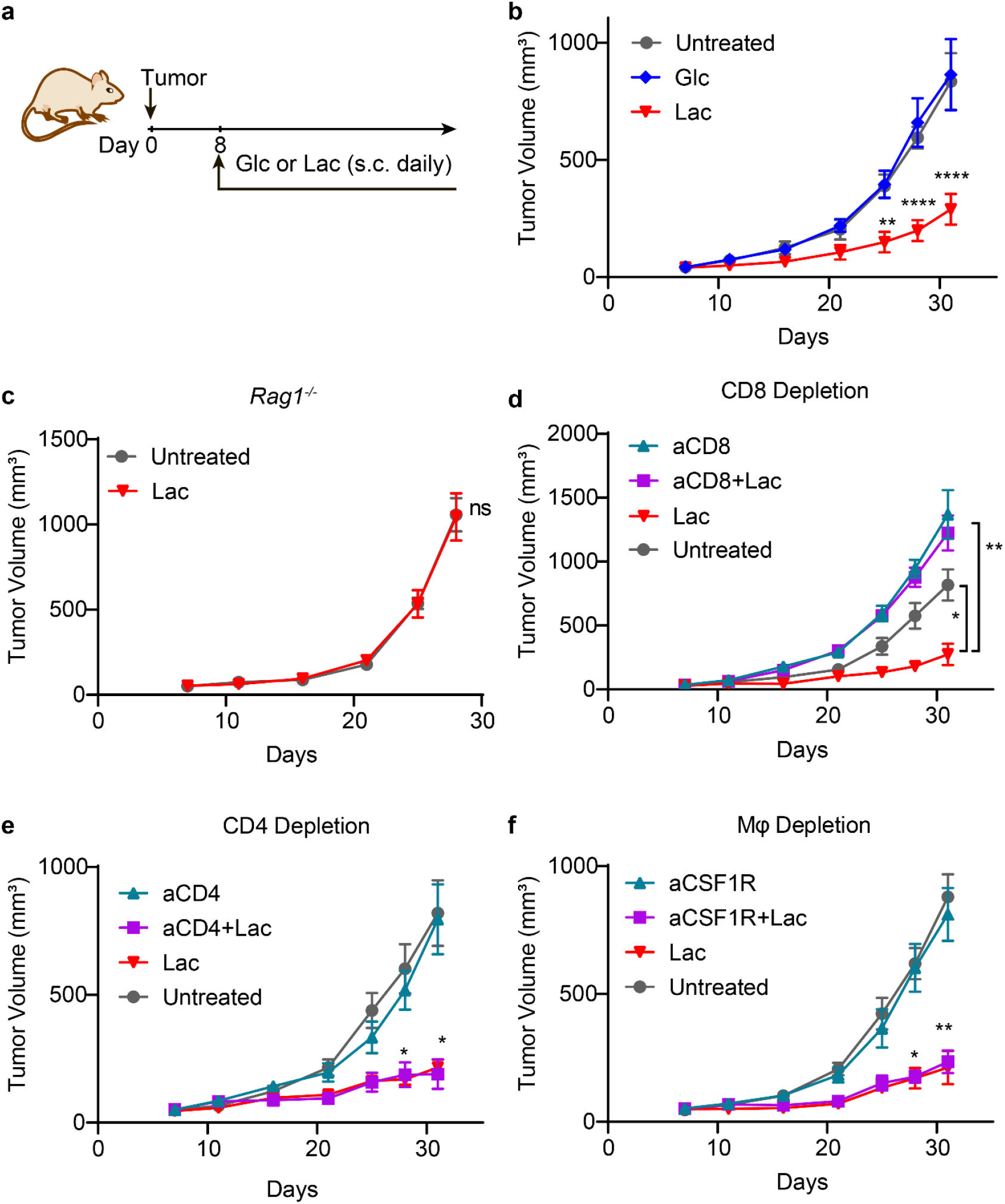
Lactate augments antitumor immunity through CD8^+^ T cells. **a**, Treatment regimen for glucose (Glc) or lactate (Lac). Glucose (5 g/kg) or lactate (1.6 g/kg) was subcutaneously administrated daily from Day 8 after tumor inoculation. **b**, Tumor growth curve of MC38 tumor model treated with glucose or lactate. C57BL/6 mice (n = 6) were inoculated with 1×10 ^6^ MC38 tumor cells and treated with glucose or lactate. **c**, Tumor growth curves of MC38 tumor in *Rag1*^-/-^ mice treated with lactate. B6.129S7-*Rag1*^*tm1Mom*^ mice (n = 7) were inoculated with 1×10 ^6^ MC38 tumor cells and treated with lactate. CD8^+^ T cell (**d**), CD4^+^ T cell (**e**) and macrophage (**f**) depletion assay in MC38 tumor model. C57BL/6 mice (n = 6) were inoculated with 1×10 ^6^ MC38 tumor cells and treated with lactate. Anti-CD8 (10 mg/kg) was administered on day 6 and then every three days until the end of the experiment. Anti-CD4 (10 mg/kg) or anti-CSF1R (20 mg/kg) was administered on day 3 and then every three days until the end of the experiment. Data are shown as means ± SEM. p value was determined by two -way ANOVA, * p < 0.05, ** p < 0.01, *** p < 0.001, **** p < 0.0001.

While lactate shows potent tumor growth inhibition in multiple animal cohorts and tumor models, no complete response was achieved with single agent treatment (**Fig. 1, Supplementary Fig. 1 a, b**). To further explore a more potent antitumor effect for mechanistic study and potential clinical application, we employed two immunotherapy regimens, which consist of anti-PD-1 as an example of checkpoint blockade therapy or PC7A nanovaccine for T cell therapy ^22^ in three murine tumor models. Glucose solution (pH 7.4, 5 g/kg) was used as a control (**Fig. 2a**).

**Fig. 2.**
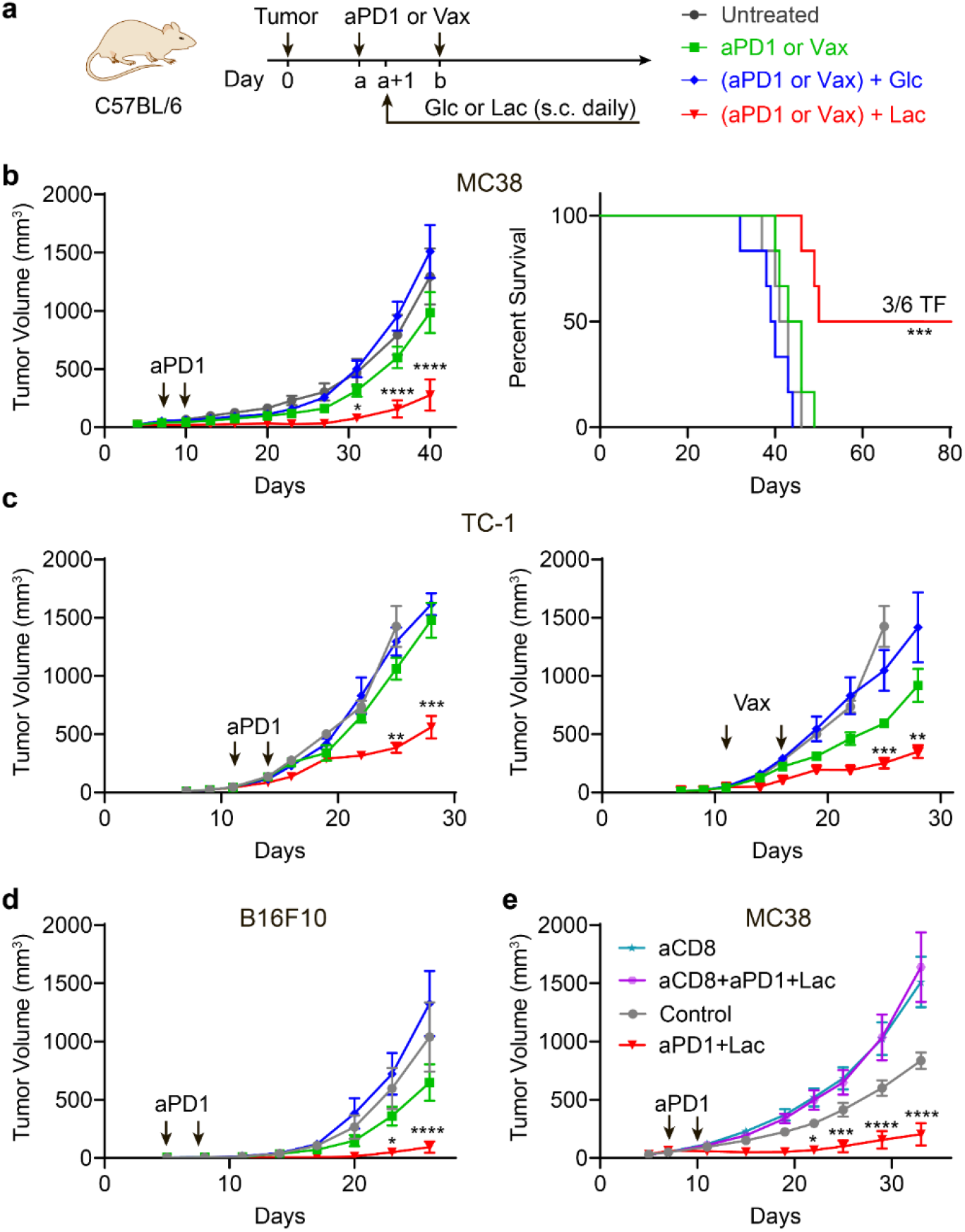
Lactate but not glucose promotes antitumor immunity in multiple tumor models. **a**, Treatment regimen for immunotherapy in combination with glucose (Glc) or lactate (Lac). Glucose (5 g/kg) or lactate (1.6 g/kg) was administrated subcutaneously daily one day after the first dose of anti-PD-1 (aPD1, i.p. injection) or PC7A vaccine (Vax, s.c. injection) treatment. **b**, Tumor growth and survival data of anti-PD-1 combined with glucose or lactate in MC38 tumor model. C57BL/6 mice (n = 6) were inoculated with 1×10^6^ MC38 tumor cells and treated with anti-PD-1 (10 mg/kg, day 7 and 10) in combination with glucose or lactate. TF: tumor free. **c**, Tumor growth curves of anti-PD-1 or PC7A vaccine combined with lactate or glucose in TC-1 tumor model. C57BL/6 mice (n = 6) were inoculated with 1.5×10 ^5^ TC-1 tumor cells and treated with anti-PD-1 (10 mg/kg, day 11 and 14) or PC7A vaccine (0.5 μg E7 peptide, day 11, 16) in combination with glucose or lactate. **d**, Tumor growth curve of anti-PD-1 combined with lactate or glucose in B16F10 tumor model. C57BL/6 mice (n = 6) were inoculated with 1.5×10 ^5^ B16F10 tumor cells and treated with anti-PD-1 (10 mg/kg, day 5 and 8) in combination with glucose or lactate. **e**, CD8^+^ T cell depletion assay in MC38 tumor model. C57BL/6 mice (n = 5) were inoculated with 1×10 ^6^ MC38 tumor cells and treated with anti-PD-1 (10 mg/kg, day 7 and 10) in combination with glucose or lactate. Anti-CD8 (10 mg/kg) was administered at day 6 and then every three days until the end of the experiment. Data are shown as means ± SEM. p value was determined by logrank test (**b**) or two-way ANOVA (**a, e–e**), * p < 0.05, ** p < 0.01, *** p < 0.001, **** p < 0.0001.

In the MC38 model, lactate treatment significantly improved the efficacy of anti-PD-1 therapy with slower tumor growth and prolonged survival (**Fig. 2b**). Combination of lactate and anti-PD-1 treatment resulted in tumor free outcomes in 50% of mice while all mice were lost in other groups before day 60. In contrast, glucose failed to improve the anti-tumor efficacy of anti-PD-1 (**Fig. 2b**). A delayed initial treatment (from day14) was also tested. Lactate shows similar anti-tumor efficacy and benefits the anti-PD1 therapy (**Supplementary Fig. 1c**). In the TC-1 tumor model which is transfected with human papilloma virus (HPV) E6/7 proteins, we observed a similar boosting effect of lactate on anti-PD-1 efficacy whereas glucose showed no effect (**Fig. 2c**). We used E7_43–62_-PC7A nanovaccine (Vax) which primes an E7-specific CD8^+^ T cell response in the TC-1 tumor model ^22^. Lactate treatment significantly improved the anti-tumor efficacy of the nanovaccine. In contrast, glucose injection abolished the anti-tumor efficacy of the nanovaccine (**Fig. 2c, Supplementary Fig. 1a**). In the B16F10 melanoma model, lactate treatment also significantly improved the efficacy of anti-PD-1 while glucose had a slight opposite effect (**Fig. 4d**). Similar to single agent therapy, the efficacy of combination therapy is dependent on CD8^+^ T cells but not on CD4^+^ T cells or macrophages (**Fig. 2e, Supplementary Fig. 1d**). Overall, these data suggest that lactate boosts anti-tumor immunity through CD8^+^T cells *in vivo*.

### Lactate treatment increases tumor infiltrating CD8^+^ T cells

To elucidate the T cell immune microenvironment of lactate-augmented anti-tumor immunity, we analyzed CD3^+^ T cells from MC38 tumors and tumor draining lymph nodes (DLNs) using single cell RNA sequencing analysis (sc-RNAseq, 10x genomics platform) after treatment by anti-PD-1 alone or combined with lactate (**Fig. 3a**). A delayed initial treatment (from day 14) was used to make sure we collect sufficient tumor tissue for downstream analysis. Overall, 21,826 single CD3^+^ T cells from both treatment conditions were subjected to the following analysis. First, we performed unsupervised graph-based clustering analysis with Seurat ^23^ to define major phenotypic T cell populations, which revealed 21 clusters and were visualized by t-distributed stochastic neighbor embedding (tSNE) algorithm (**Fig. 3b, Supplementary Fig. 2**). Using well-established marker genes (e.g., *Cd8a, Pdcd1, Cd4, Foxp3, Ctla4, etc*.), we identified 11 CD8^+^ T cell, 7 CD4^+^ helper T cell (CD4^+^ Th), and 3 CD4^+^ regulatory T cell (CD4^+^ Treg) clusters (**Fig. 3 b, c, Supplementary Fig. 2**).

**Fig. 3.**
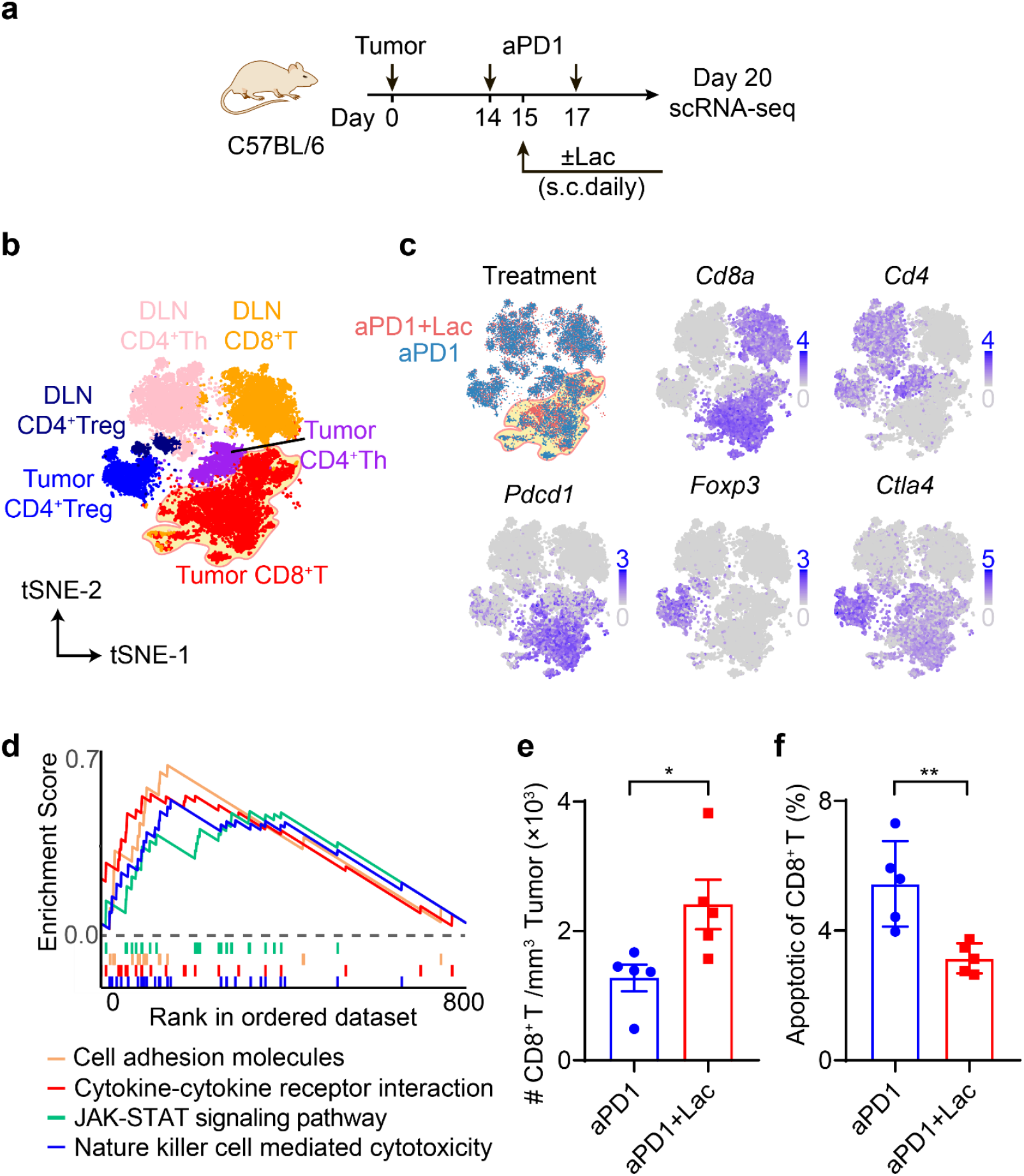
Lactate treatment increases infiltrating CD8^+^ T cells in MC38 tumors. **a**, Experimental design of single cell transcriptomic analysis of anti-PD-1 with or without lactate. C57BL/6 mice were inoculated with 1×10^6^ MC38 tumor cells and treated with anti-PD-1 (10 mg/kg, day 14 and 17) and lactate (1.6 g/kg, s.c. daily from day 15 to 19). Tumor and tumor draining lymph nodes were harvested on day 20 and analyzed by single cell RNA sequencing using the 10x platform. **b**, tSNE plot of T cell clusters with location and cell type information analyzed with Seurat v3.0.1. DLN: tumor draining lymph nodes. **c**, Distribution of T cells from different treatments and expression of marker genes. **d**, Significantly upregulated pathways in tumor infiltrating CD8^+^ T cells after lactate treatment by unbiased gene set enrichment analysis (gene set database: c2.cp.kegg.v7.2.symbols). **e**, Validation of increased tumor infiltrating CD8^+^ T cells by flow cytometry. C57BL/6 mice (n = 5) were inoculated with 1×10^6^ MC38 tumor cells and treated with anti-PD-1 (10 mg/kg, day 14 and 17) and lactate (1.6 g/kg, s.c. daily from day 15 to 19). MC38 tumors were harvested on day 20 and analyzed by flow cytometry. **f**, Analysis of apoptosis markers of CD8^+^ T cells in tumor microenvironment by flow cytometry. Same samples as described in E. Data are shown as means ± SEM. p value was determined by t test (**e-f**), * p < 0.05, ** p < 0.01.

Lactate treatment increased the total number of tumor infiltrating CD8^+^ T cells (**Fig. 3c**). Gene set enrichment analysis of tumor infiltrating CD8^+^ T cells showed that lactate treatment induced a significant upregulation of T cell function and signaling related genes (e.g., *Fasl, Gzmb, Prf1, Il2ra, Ifngr1, Il*7*r* and *Ccl5*) and pathways (e.g., JAK-STAT and cytokine-cytokine receptor interaction, **Fig. 3d, Supplementary Fig. 3**). Flow cytometry analysis validated the increased CD8^+^ T cell infiltration in MC38 tumors following lactate treatment (**Fig. 3e**). Furthermore, lactate treatment significantly reduced the percentage of apoptotic CD8^+^ T cells as gated by active Caspase 3 (**Fig. 3f**).

The 11 CD8^+^ T cell clusters follow a typical differentiation process from naïve to exhausted states observed during anti-tumor immune response (**Supplementary Fig. 2**, see **Data_file_S1** for marker gene list). Among them, we identified 2 naïve T cell clusters (CD8-1 and CD8-2), 2 stem-like (pre-exhausted) T cell clusters (CD8-5 and CD8-6), 4 exhausted T cell clusters (CD8-7 through CD8-10) and 3 cell clusters in intermediate status (CD8-3, CD8-4 and CD8-11). Chi-square analysis (shown as ratio of observed to expected cell numbers, R_O/E_) revealed that lactate treatment led to a consistent increase in cell numbers in stem-like T cell clusters in tumor while exhausted clusters responded differentially (**Supplementary Fig. 2**).

### Lactate treatment increases stem-like CD8^+^ T cell population in MC38 tumors

To further investigate the effect of lactate on different CD8^+^ T cell subtypes, we analyzed the differentiation trajectory of tumor infiltrating CD8^+^ T cells using pseudotime analysis by Monocle 2 (**Fig. 4a, Supplementary Fig. 4a**) ^24^. The trajectory was divided into 7 states with unsupervised clustering, which begins with a stem-like state 1 and differentiates into exhausted states. Stem-like CD8^+^ T cells in state 1 have high abundance of stem-like genes (*Tcf7, Il7r, Cxcr3*) and low abundance of effector genes (*Ifng, Gzmb, Gzmc, Gzmf*) and exhaustion genes (*Lag3, Pdcd1, Havcr2, Entpd1*) (**Fig. 4b, Fig. S4b** and **Data_file_S2**).

**Fig. 4.**
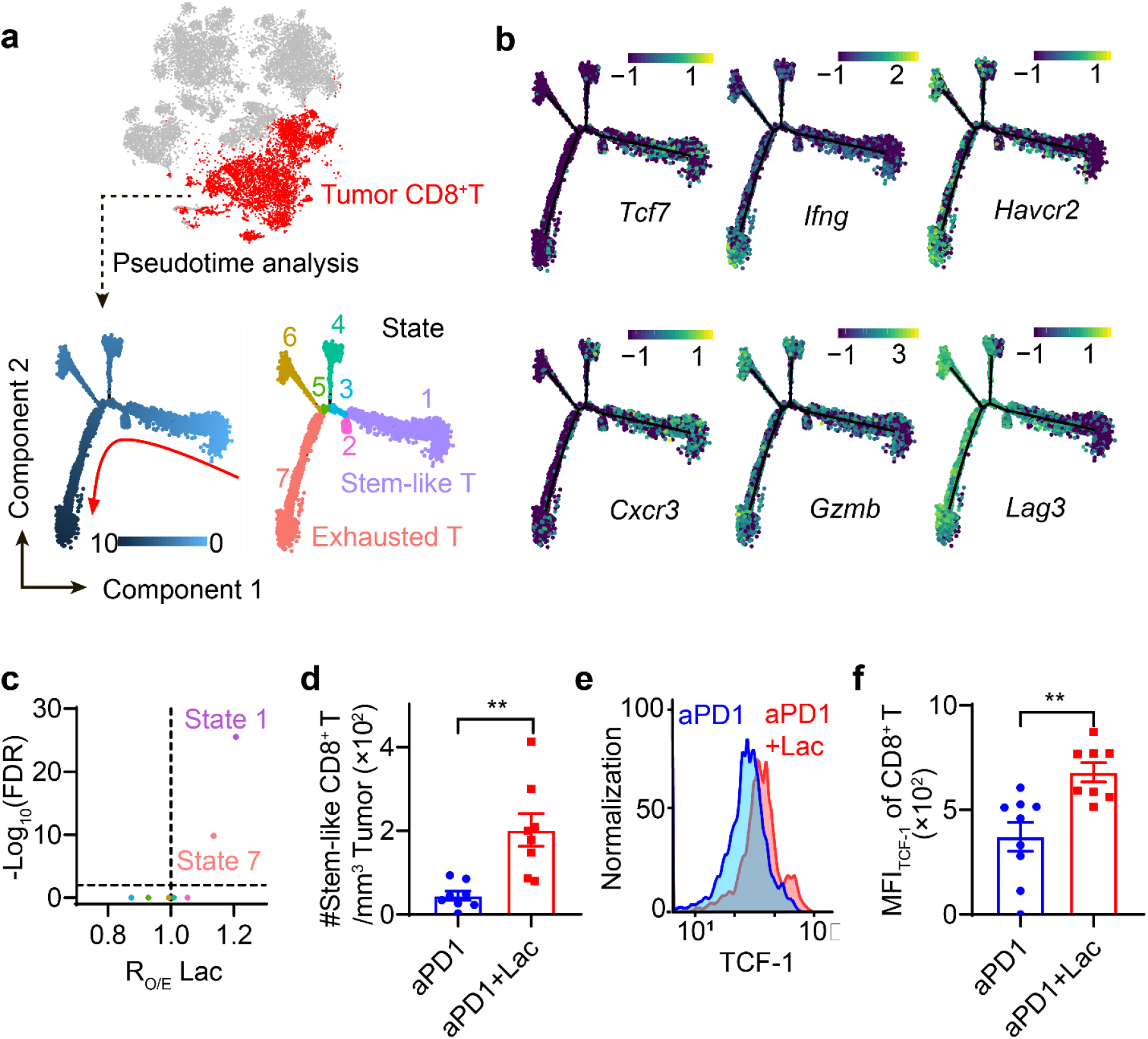
Lactate treatment increases stem-like CD8^+^ T cell population in MC38 tumors. **a**, Pseudotime trajectory of CD8^+^ T cells in tumors identifies the differentiation process and distinct states of CD8^+^ T cell subtypes. **b**, Labeling of top marker genes on pseudotime trajectory identifies the cells in state1 are stem-like T cells while those in state 7 are exhausted T cells. **c**, Change of CD8^+^ T cell fraction with aPD1+Lac verses aPD1 treatment. R_O/E_Lac analysis showed lactate treatment increased the T cell populations in states 1 and 7. **d**, Validation of increase in TCF1^+^ PD1^+^ Lag3^-^ CD8^+^ T cell population after lactate treatment by flow cytometry. C57BL/6 mice (n = 8) were inoculated with 1×10 ^6^ MC38 tumor cells and treated with anti-PD-1 (10 mg/kg, day 14 and 17) and lactate (1.6 g/kg, s.c. daily from day 15 to 19). Tumors were harvested on day 20 and analyzed by flow cytometry. **e-f**, Flow cytometry plot and quantification of TCF-1 expressions in tumor-infiltrating CD8^+^ T cells after different treatments. Samples are the same as described in D. Data are shown as means ± SEM. p value was determined by Chi -square test (**c**) or t test (**e, f**), * p < 0.05, ** p < 0.01.

We observed that lactate treatment resulted in the most significant increase in the number of stem-like CD8^+^ T cells (state 1) over all other T cell states (**Fig. 4c**). We used flow cytometry to confirm the increase in the number of tumor infiltrating stem-like T cells (**Fig. 4d**) and elevated expression of the TCF-1 protein in the total population of tumor infiltrating CD8^+^ T cells (**Fig. 4 e,f**) in response to lactate treatment. Collectively, these results show that lactate treatment increased the number of stem-like T cells and elevated the expression of *Tcf7*/TCF-1 in CD8^+^ T cells *in vivo*.

### Lactate increases TCF-1 expression and reduces apoptosis of CD8^+^ T cells during *ex vivo* expansion

To elucidate the effect of lactate on CD8^+^ T cells, we first investigated whether lactate impacts the *ex vivo* culture of CD8^+^ T cells derived from mouse splenocytes or human peripheral blood mononuclear cells (PBMCs). OT-I splenocytes were obtained from C57BL/6-Tg(TcraTcrb)1100Mjb/J (OT-I transgenic) mice and stimulated with OVA peptide (OVA_p_, SIINFEKL) and IL-2 for two days, followed by cell culture with anti-CD3, anti-CD28 and IL-2 (**Fig. 5a**). The dose of lactate for *in vitro* T cell culture was set to 40 mM to mimic the lactate concentration in tumor interstitial fluid (TIF) (**Supplementary Fig. 5a**). Lactate treatment significantly increased the number of TCF-1^hi^CXCR3^hi^ cells and mean fluorescent intensity of TCF-1 in CD8^+^ T cells compared to control condition (RPMI medium containing 2 mM lactate, **Fig. 5 b,c** and **Supplementary 5b**). Similar to our *in vivo* observation (**Fig. 3f**), lactate treatment during *ex vivo* expansion reduced the percentage of apoptotic CD8^+^ T cells (**Fig. 5d** and **5e**). Of note, by examining the co-staining of TCF-1 and active caspase 3, apoptotic CD8^+^ T cells were enriched in the low TCF-1 expressing population (**Fig. 5d**). This result is consistent with reports of *Bcl2* induction by *Tcf7* upregulation, which inhibits apoptosis ^25-27^. Finally, we quantified the mRNA expressions for genes of interest during the T cell expansion by real-time polymerase chain reaction (PCR). Lactate treatment significantly increased the expression level of *Tcf7* mRNA, while insignificant differences were observed with activation/exhaustion related genes including *Pdcd1, Lag3* and *Ifng* and metabolism related genes including *Acss1, Acss2, Hif1a, Ldha and Ldhb* (**Fig. 5f**).

**Fig. 5.**
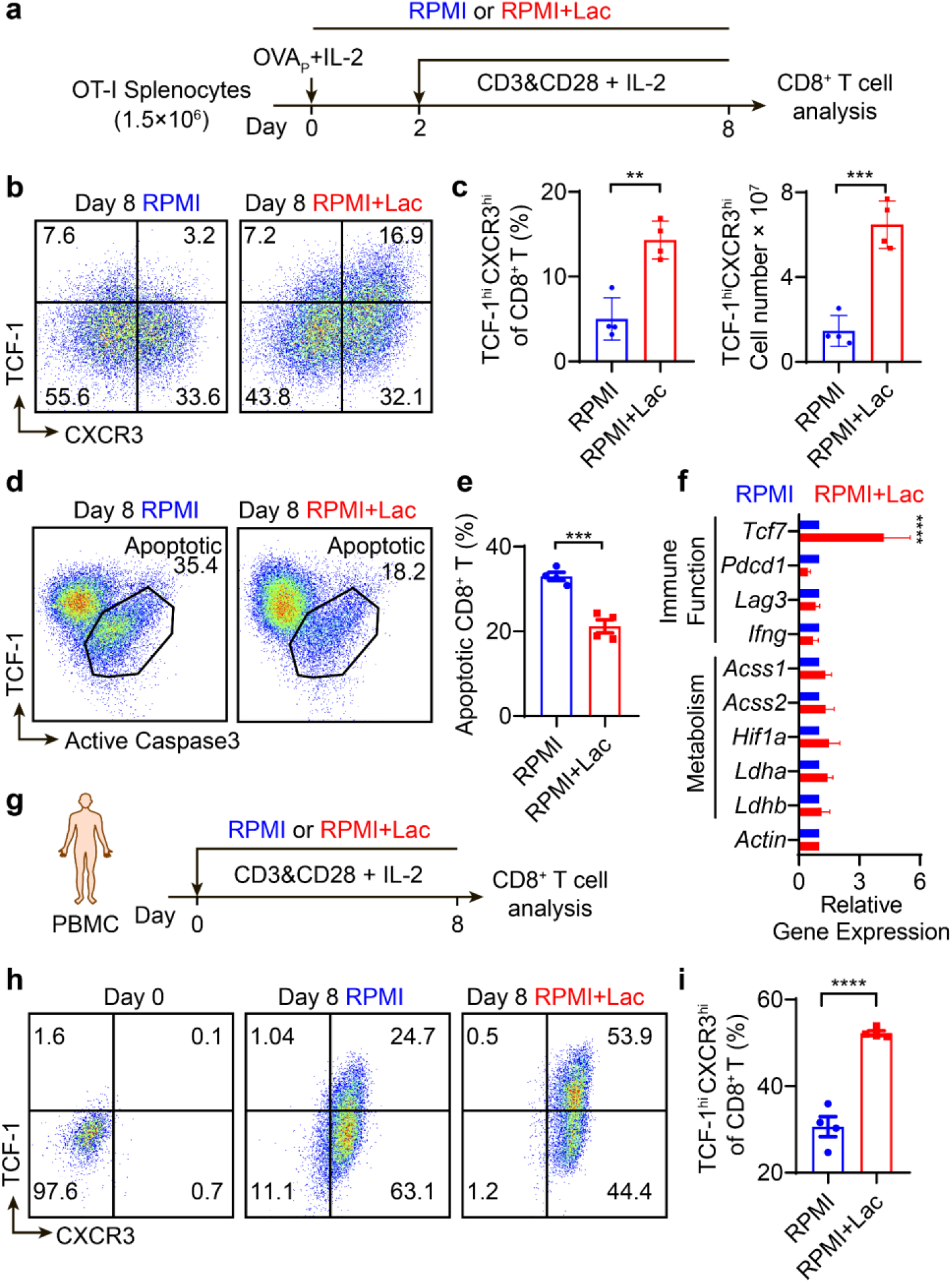
Lactate increases TCF-1 expression and reduces apoptosis of CD8^+^ T cells during *ex vivo* expansion. **a**, Experimental design of *ex vivo* OT-I CD8^+^ T cell expansion. Fresh splenocytes from C57BL/6-Tg(TcraTcrb)1100Mjb/J mice were primed with SIINFEKL peptide (1 μg/mL) and hIL-2 (50 U/mL) for two days and stimulated with anti-CD3 and anti-CD28 (0.5 μg/mL each) from day 3 to 8 with hIL-2 (30 U/mL). **b-c**, Flow cytometry plots and quantification of TCF-1^hi^CXCR3^hi^ population of OT-I CD8^+^ T cells on day 8 of *ex vivo* expansion. **d-e**, Quantification of percentage of apoptotic OT-I CD8^+^ T cells by flow cytometry on day 8 of *ex vivo* expansion. **f**, Relative gene expression in OT-I CD8^+^ T cells on day 4 detected by RT-PCR. **g**, Experimental design of *ex vivo* expansion of CD8^+^ T cells from human PBMCs. PBMCs from cord blood were activated and cultured in the presence of anti-CD3 and anti-CD28 beads (T cell : Beads =1 : 1) supplemented with hIL-2 (30 U/mL). **h-i**, Flow cytometry plots and quantification of TCF-1^hi^CXCR3^hi^ population of human CD8^+^ T cells on day 8 of *ex vivo* expansion. Data are shown as means ± SEM. p value was determined by t test, ** p < 0.01, *** p < 0.001, ****p < 0.0001.

To study the lactate effect on human CD8^+^ T cells, we employed PBMCs isolated from human cord blood. PBMCs were cultured in lactate-augmented (40 mM) or control RPMI medium with anti-CD3 and anti-CD28 modified microbeads and IL-2 for 8 days (**Fig. 5g**). Similar to the results from OT-I CD8^+^ T cells, lactate treatment significantly increased the TCF-1^hi^CXCR3^hi^ cell population and the mean florescent intensity of TCF-1 protein in human CD8^+^ T cells (**Fig. 5h, i and Supplementary Fig. 5c**). In contrast, lactic acid, the conjugated acid form of lactate, resulted in significant CD8^+^ T cells death *in vitro* (**Supplementary Fig. 5d**).

### Lactate induces T cell stemness through epigenetic regulation

Besides its newly uncovered role as a primary carbon fuel source, lactate has additional functions as an agonist to G-protein-coupled receptor (GPCR) signaling and as an inhibitor to histone deacetylase (HDAC). As a carbon fuel source, lactate is converted to pyruvate by lactate dehydrogenase B and further fuels the TCA cycle ^16, 17^. As a GPCR agonist (e.g., GPR81), lactate regulates multiple biological processes through the secondary messenger cyclic AMP (cAMP) ^28, 29^. Also, lactate has been reported as an HDAC inhibitor with an IC_50_ of 40 mM ^30^. The increase of TCF-1 expression in CD8^+^ T cells may arise from lactate augmented metabolites (e.g., pyruvate, amino acids), lactate-mediated GPCR signaling or HDAC inhibition (**Fig. 6a**).

**Fig. 6.**
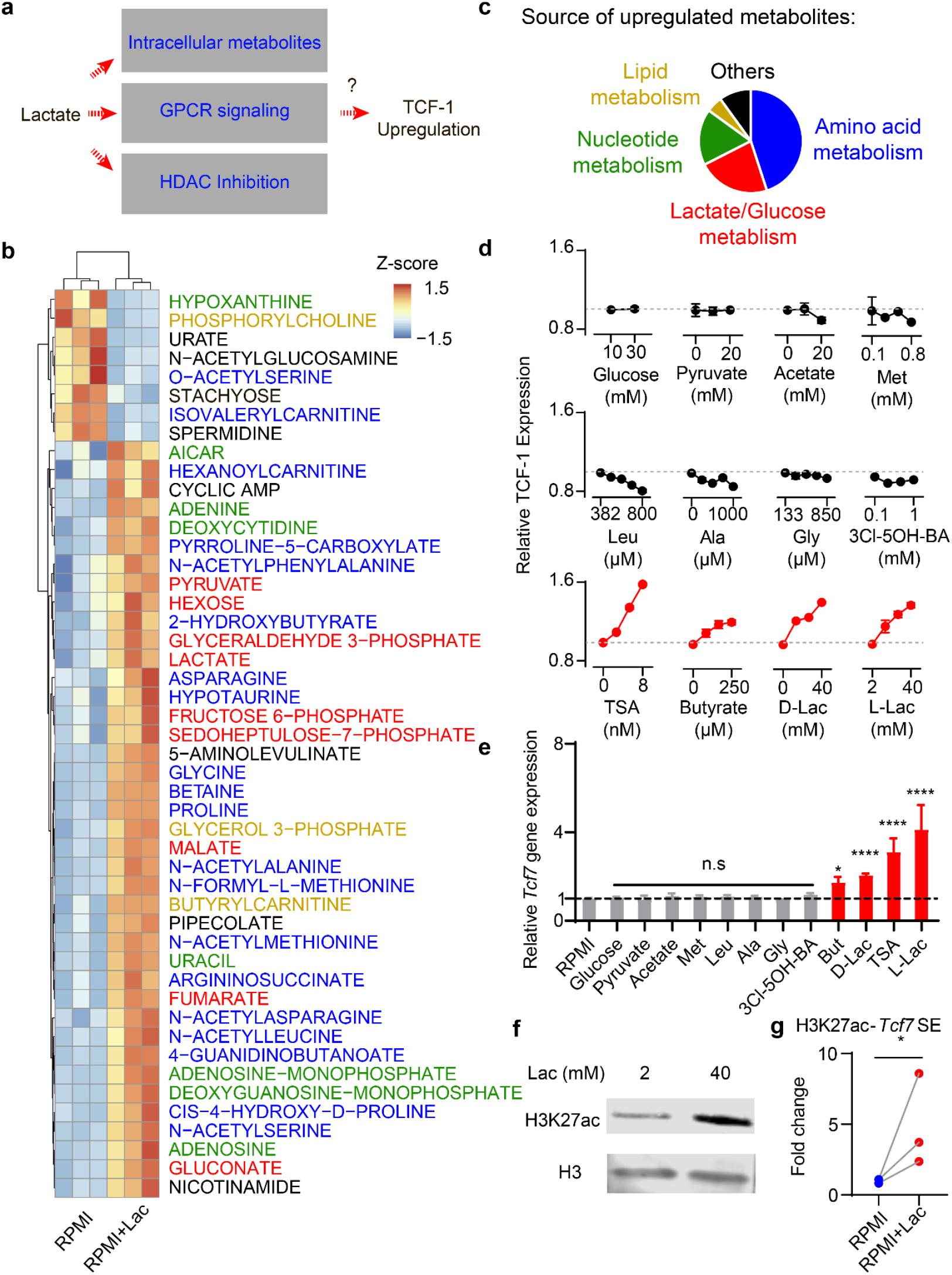
Lactate increases TCF-1 expression through inhibition of histone deacetylases (HDAC). **a**, Potential mechanisms for lactate induced TCF-1 upregulation. **b**, Heatmap of significantly changed metabolites in CD8^+^ T cells treated with or without 40 mM lactate. **c**, Source of upregulated metabolites with 40 mM lactate treatment. **d**, TCF-1 protein expression in CD8^+^ T cells treated by different metabolites, GPCR agonist or HDAC inhibitors. **e**, Gene expression of *Tcf7* in CD8^+^ T cells treated by different metabolites, GPCR agonist or HDAC inhibitors. **f**, Western blot of histone H3K27ac of CD8^+^ T cells cultured with or without 40 mM lactate. **g**, Quantification of H3K27ac enrichment by CUT&RUN PCR at the *Tcf7* super enhancer locus. Data are shown as means ± SEM. p value was determined by t –test (**d**) or ratio paired t test (**f**), * p<0.05, ****P <0.0001.

To investigate the biochemical mechanism, we first measured the abundance of ∼200 common metabolites in CD8^+^ T cells (OT-I) cultured with lactate-augmented (40 mM) or control RPMI medium (**Fig. 6b**). Lactate treatment boosted central carbon metabolism, including a variety of metabolites in glucose/lactate, amino acid, nucleotide and lipid metabolisms (**Fig. 6b,c** and **Data_file_S3**). The increased abundance of lactate, pyruvate, TCA cycle intermediates and multiple amino acids correlates with lactate influx, conversion to pyruvate and subsequent utilization in the TCA cycle as shown by the stable isotope tracing study (**Supplementary Fig. 6, Data_file_S4**). In regards to the GPCR signaling pathway, lactate treatment increased cAMP levels in the T cells.

We next examined the expression of TCF-1 in the CD8^+^ T cells as a function of selected metabolites or a GPR81 agonist (**Fig. 6 d,e**). Increasing glucose concentration from 10 to 25 mM in RPMI medium showed no effect on TCF-1 expression. Pyruvate (0-20 mM), the direct metabolite downstream of lactate conversion in cells, did not cause increase of TCF-1. Acetate, another monocarboxylate species that can be synthesized from pyruvate and fuel TCA cycle through acetyl-CoA, also had little effect. Lactate augmented amino acids, including essential amino acids (e.g., Methionine and Leucine) and nonessential amino acids (e.g., Glycine and Alanine), also did not increase TCF-1 expression. A GPR81 ligand, 3-chloro-5-hydroxybenzoic acid (3Cl-5OH-BA), which stimulates cAMP production similar to lactate-induced effect ^29, 31^, didn’t increase TCF-1. These results indicate lactate-mediated HDAC inhibition as a potential driver for increased T cell stemness.

Lactate has been reported as an endogenous inhibitor of HDAC that links the metabolic state of cells to gene transcription ^30^. To further corroborate the role of HDAC inhibition on T cell stemness, OT-I CD8^+^ T cells were treated with known HDAC inhibitors including trichostatin A (TSA), D-lactate (all other lactates refer to L-lactate unless noted otherwise), or butyrate. Flow cytometry and qPCR analyses showed elevated TCF-1 expressions similar to the treatment outcome by 40 mM lactate (**Fig. 6d,e**). Epigenomic profiling identified acetylation of H3K27 at the *Tcf7* super enhancer site correlates with the naïve and central memory identity of CD8^+^ T cells ^32^. Western blot of lactate treated CD8^+^ T cells showed 3.5-fold increase of acetylation at the H3K27 site (**Fig. 2f**). Cleavage under targets and release using nuclease (CUT&RUN) PCR analysis further confirmed significant H3K27ac enrichment at the *Tcf7* super enhance locus in the lactate treated CD8^+^ T cells (**Fig. 6g**).

Collectively, these results indicate that lactate directly induces CD8^+^ T cell stemness through epigenetic regulation of cell identity.

### Adoptive transfer of lactate-pretreated CD8^+^ T cells achieves potent tumor growth inhibition *in vivo*

Recent studies have linked TCF-1 expressing stem-like CD8^+^ T cells, which retain polyfunctionality and persistence of cytotoxic function, as the key T cell subsets which respond to adoptive cell transfer therapy, anti-PD-1 and vaccination therapy^33-39^. To investigate whether lactate-induced TCF-1 expressing CD8^+^ T cells can improve tumor therapy, we harvested and intravenously injected OVA-specific CD8^+^ T cells into MC38-OVA tumor bearing mice (**Fig. 7a**). OVA-specific T cells pretreated with lactate *ex vivo* demonstrated significantly improved tumor growth inhibition over those cultured in control RPMI medium (**Fig. 7b**). To further investigate the effect of transferred vs. endogenous CD8^+^ T cells on antitumor efficacy, we employed OVA tetramer^+^ CD8^+^ T cells from CD45.2 expressing donors and transferred them to CD45.1 expressing tumor bearing mice (**Fig. 7c**). Seven days after cell transfer, we analyzed the number of host (CD45.1^+^) and donor (CD45.2^+^) CD8^+^ T cells in tumors by flow cytometry. Lactate pretreatment significantly expanded CD45.2^+^ tetramer^+^ CD8^+^ T cells to a larger clonal size over those from control RPMI medium (**Fig. 7 d,e**). In contrast, the number of endogenous CD45.1^+^ tetramer^+^ CD8^+^ T cells showed no significant difference after transfer of CD45.2^+^ T cells with and without lactate pretreatment (**Fig. 7f**).

**Fig. 7.**
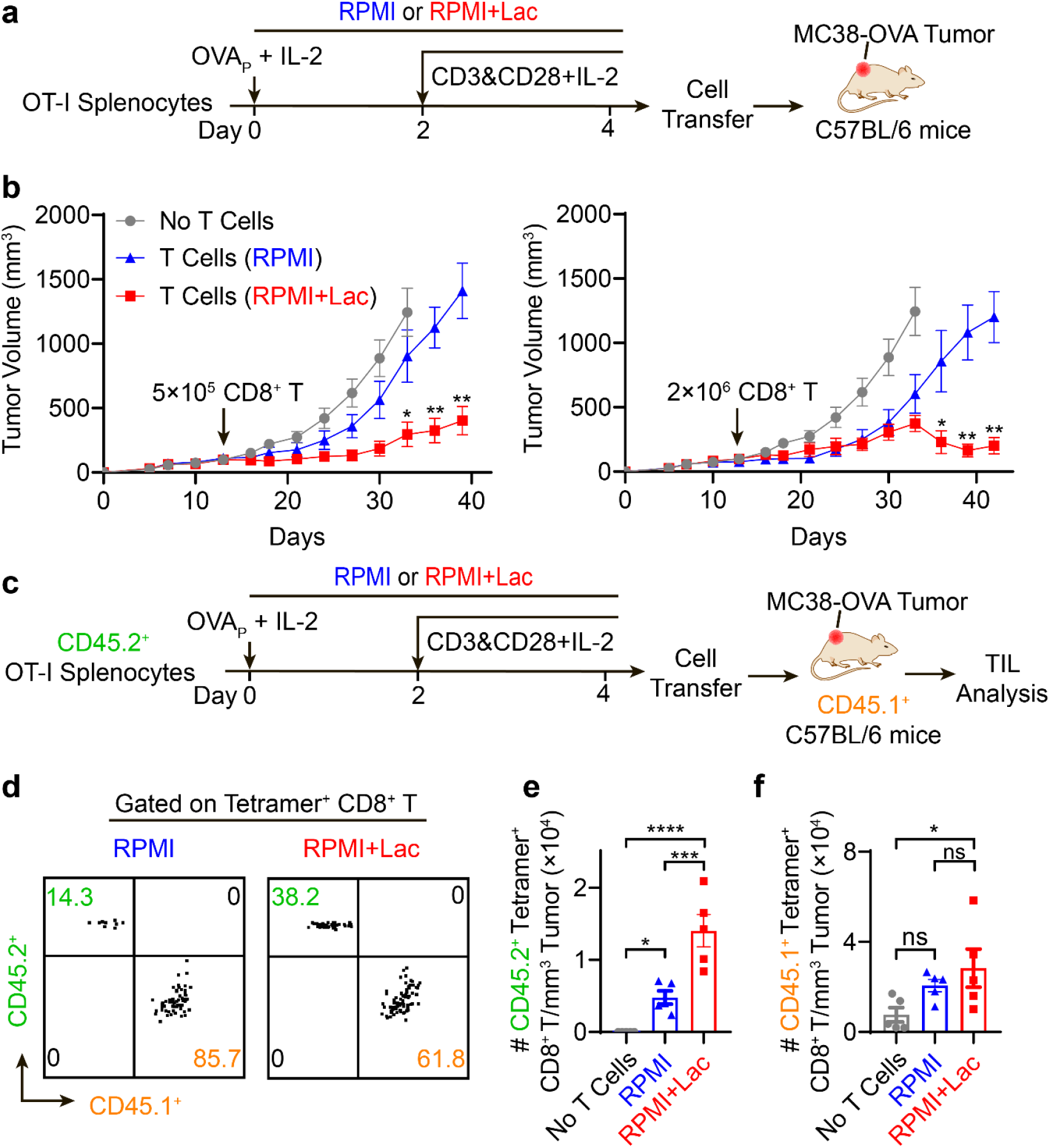
CD8^+^ T cells expanded under high lactate condition show potent tumor growth inhibition *in vivo*. **a**, Treatment plan of T cell receptor engineered T cell therapy (TCR-T). OT-I splenocytes were cultured *ex vivo* for 4 days with or without 40 mM sodium lactate in the culture medium. After purification with negative selection magnetic beads, CD8^+^ T cells were transferred to MC38-OVA tumor bearing mice. Average tumor size is above 100 mm^3^ at the time of cell transfer. **b**, The growth curves of MC38-OVA tumor model were significantly inhibited after transfer of TCR-T (5×10 ^5^ or 2 ×10 ^6^) pretreated with sodium lactate. **c**, Analysis of tumor-infiltrating T cells after adoptive TCR-T cell transfer. Seven days after the cell transfer, tumor-infiltrating lymphocytes were analyzed by flow cytometry. **d-f**, Flow cytometry plot and quantification of transferred (CD45.2^+^) and endogenous (CD45.1^+^) OVA-tetramer^+^ CD8^+^ T cells show lactate pretreatment significantly increased the number of transferred CD8^+^ T cells (CD45.2^+^) but not endogenous CD8^+^ T cells (CD45.1^+^) in the MC38-OVA tumors. Data are shown as means ± SEM. p value was determined by two -way ANOVA (**b, c**) or one-way ANOVA (**e, f**), ns no significance, * p<0.05, ** p<0.01, *** p<0.001, ****p<0.0001.

These data support that lactate-pretreated CD8^+^ T cells, which have high TCF-1 expression, show improved anti-tumor immune response *in vivo*.

## Discussion

In this study, we report a mechanistic link between lactate, CD8^+^ T cell stemness and improved outcome of cancer immunotherapy. Recent studies show lactate overtakes glucose as the primary fuel source of most tissues ^17, 18^. Our side-by-side comparison between subcutaneous administration of high dose lactate versus glucose revealed a surprising immune protective role of lactate (**Fig. 1-2**). We hypothesize the immune protective role of lactate is undermined by its conjugated acid form, lactic acid. In the highly glycolytic and poorly vascularized microenvironment of solid tumors, glucose is preferentially converted to lactic acids (aka Warburg effect) by cancer cells or myeloid cells, which are further secreted to the extracellular environment through monocarboxylate transporters ^40-42^. Multiple studies have shown lactic acid is responsible for retardation of immune cell functions leading to tumor immune evasion ^19-21, 43^. Our data also showed lactic acid mediated significant CD8^+^ T cell death in culture, whereas lactate did not (**Supplementary Fig. 5d**). In contrary, lactate elevated the stemness of CD8^+^ T cells *ex vivo* and *in vivo* which leads to immune protection against cancer.

Lactate has broad physiological functions in carbon metabolism, cell signaling and epigenetic regulation ^17, 28^-^30, 44^. Our mechanistic studies reveal lactate-mediated HDAC inhibition is primarily responsible for increased stemness of CD8^+^ T cells. In contrast, glucose, lactate-augmented metabolites (e.g., pyruvate, various amino acids) or GPCR signaling (e.g., GPR81 agonist) did not appear to contribute to the T cell stemness. Known HDAC inhibitors (e.g., TSA, butyrate) also increased T cell stemness in the current experimental conditions (**Fig. 6d**). However, prolonged exposure at elevated concentrations of these agents resulted in decreased viability of T cells (data not shown). In contrast, *ex vivo* expansion of CD8^+^ T cells under high lactate concentrations reduced cell apoptosis over prolonged culture (**Fig. 5**). As an endogenous carbon fuel source, lactate offers a wider therapeutic window with minimal cell toxicity. These results add to the growing evidence in the metabolic control of immune functions by nutrients or metabolites ^1-3^.

As a common metabolite, lactate has broad influence on many different cell types (e.g. T cells, cancer cells, macrophages, etc.) in the tumor microenvironment. A recent study shows LDHA-mediated lactic acid fuels CD4^+^ regulatory T cells *(21)*. Our study provides an initial observation of the lactate effect on stem-like CD8^+^ T cell and shows that lactate benefits anti-tumor immune response instead of being immune suppressive. With the cell depletion assay (**Fig 1c-e, 2e, Supplementary Fig. 1d**) and adoptive transfer of lactate-cultured CD8^+^ T cells (**Fig. 7**), we conclude that the boosting of CD8^+^ T cell stemness is a sufficient condition for the anti-tumor efficacy of lactate. Meanwhile, the effect of lactate on cancer cells, tumor acidity and global immunity will need further investigation.

In summary, results from this study elucidated the role and mechanism of lactate in antitumor immunity. Single cell transcriptomics and flow cytometry analysis revealed an increased subpopulation of stem-like TCF-1-expressing CD8^+^ T cells upon lactate treatment, which is further confirmed by *ex vivo* culture of CD8^+^ T cells from mouse splenocytes and human PBMCs. Inhibition of histone deacetylase activity by lactate is responsible for the increased TCF-1 expression on the CD8^+^ T cells. Lactate has the potential to augment the therapeutic outcomes of immune checkpoint blockade, T cell vaccine and adoptive T cell transfer therapy.

## Methods Reagents

The details of reagents used are provide in Table S1.

### Mice

Female C57BL/6J, NOD.Cg-Prkdc^scid^/J C57BL/6-Tg(TcraTcrb)1100Mjb/J, B6.129S7-*Rag1*^*tm1Mom*^/J, B6.SJL-Ptprc^a^Pepc^b^/BoyJ mice were purchased from Jackson Laboratory. All mice were maintained in specific pathogen free animal facility. All animal experiments were performed with ethical compliance and approval from Institutional Animal Care and Use Committee of the University of Texas Southwestern Medical Center.

### Cells

Murine melanoma B16F10 and colon adenocarcinoma MC38 cell lines were purchased from American Type Culture Collection (ATCC). TC-1 cells were kindly provided by Dr. T. C. Wu at John Hopkins University. MC38-OVA cells were made by lentiviral transduction of OVA gene. All cancer cell lines were routinely tested using e-Myco mycoplasma PCR Detection kit (Bulldog Bio Inc) and cultured in high glucose Dulbecco’s modified Eagle’s medium supplemented with 10% fetal bovine serum, 100 U/mL penicillin, 100 U/mL streptomycin and 1×GlutaMax under 5% CO_2_ at 37 °C. Immune cells from mouse or human were cultured in RPMI 1640 medium supplied with 10% heat-inactivated fetal bovine serum, 100 U/mL penicillin, 100 μg/mL streptomycin, 20 mM HEPES, 50μM 2-Mercaptoethanol and 1×GlutaMax.

### Human Peripheral Blood Mononuclear Cells

Human cord blood samples were obtained from UT Southwestern (UTSW) Parkland Hospital according to the regulation and the use approval of human cord blood at UTSW medical center. Human peripheral blood mononuclear cells (PBMC) were purified from cord blood by Ficoll-Paque Plus according to the manufacturer’s manual.

### Tumor Growth and Treatment

C57BL/6J mice were inoculated with 1×10 ^6^ MC38 tumor cells, or 1.5×10 ^5^ TC-1 tumor cells or 1.5×10 ^5^ B16F10 tumor cells on the right flank on day 0. For MC38 model, animals were intraperitoneally (i.p.) treated with anti-PD-1 (10 mg/kg, day 7 and 10) in combination with glucose or lactate (s.c., 5 g/kg or 1.6 g/kg, respectively) daily, beginning on day 8. For TC-1 tumor model, animals were treated with anti-PD-1 (i.p. 10 mg/kg, day 11 and 14) or PC7A vaccine (s.c. 0.5 μg E7 peptide, day 11 and 16) in combination with glucose or lactate (s.c., 5 g/kg or 1.6 g/kg, respectively) daily, beginning on day 12. For B16F10 tumor model, animals were treated with anti-PD-1 (i.p. 10 mg/kg, day 5 and 8) in combination with glucose or lactate (s.c., 5 g/kg or 1.6 g/kg, respectively) daily, beginning on day 6. For immune cell depletion assay, anti-CD8 antibodies, anti-CD4 antibodies or anti-NK1.1 were administrated every three days during the treatment (i.p. 10 mg/kg). For single cell analysis and *in vivo* flow cytometry analysis in MC38 tumor model, animals were intraperitoneally (i.p.) treated with anti-PD-1 (10 mg/kg, day 14 and 17) in combination with or without lactate (s.c., 1.6 g/kg) daily, beginning on day 15. Tumor and tumor draining lymph nodes were collected on day 20 for analysis. Tumor volumes were measured with a caliper by the length (L), width (W) and height (H) and calculated as tumor volume = L×W×H/2.

### Lactate Concentration in tumor interstitial fluid

Tumor interstitial fluid was collected from freshly resected MC38 tumor. Specimens were centrifuged against a 70 μm cell strainer at 4 °C for 5 min at 300g. Flow -through tissue interstitial fluid was centrifuged at 4 °C for 5 min at 500g. Supernatant were flash-frozen and stored at −80 °C before batch analysis. The lactate concentration was determined with lactate assay kit (sigma-aldrich, MAK064) according to the manufacturer’s protocol.

### Single Cell RNA Sequencing

Tumor-infiltrating CD3^+^ T cells were obtained by microbeads enrichment and flow sorting. Briefly, CD45^+^ cells from tumors were isolated with Tumor Dissociation Kit (Miltenyi) and purified with Dead Cell Removal Kit (Miltenyi) and CD45 (TIL) MicroBeads (Miltenyi) according to the manufacturer’s instructions. The purified cells were further sorted for live CD45^+^CD3^+^ population on BD Aria II SORP sorter. Tumor draining lymph node cells were flow sorted for live CD45^+^ population. All collected cells were barcoded with Chromium Single Cell 5’ Library & Gel Bead Kit (10x Genomics). Library construction was performed following the protocol of Chromium Single Cell 5’ Library Construction Kit. Libraries were pooled with Chromium i7 Multiplex Kit. The final pooled libraries were sequenced on NovaSeq S4 platform (Novogene).

### Analysis of Single Cell RNA Sequencing Data

Sequencing reads were aligned to mouse genome (mm10) and unique molecular identifiers (UMIs) were counted for each gene using Cell Ranger 3.0.1 (10x Genomics) The R software package Seurat (version 3.0.1) was used for further analysis. Cells with a high or low proportion of detected genes (> 2 × median absolute deviations) and those with a high of mitochondrial (> 15%) were removed. Cells were divided into clusters with graph-based clustering using the Seurat function Find Clusters at resolution 0.8. Marker genes for each cell cluster were calculated with Wilcoxon Rank Sum test with log10 (fold change) >0.1. Monocle 2 (version 2.10.1) was used to estimate a pseudo-temporal path of tumor infiltratingCD8^+^ T cell differentiation. Monocle object was formed by Monocle implemented newCellDataSet function with lowerDetectionLimit = 0.5. Differentially expressed genes (DEGs) were identified by limma R package with an absolute log2 fold change (FC) > 0.05 and q < 0.01 Pathway enrichment was performed on ranked DEG lists with GSEA using KEGG under gene size 15 and FDR 0.05.

### *Ex Vivo* Culture of Activated CD8^+^ T Cells

For mouse studies, single cell suspensions from lymph nodes and spleens of C57BL/6-Tg (TcraTcrb)1100Mjb/J transgenic mice were activated with SIINFEKL peptide (1 μg/mL) and hIL-2 (50 IU/mL). 48 hr post activation, CD8^+^ T cells were further cultured with anti-CD3 and anti-CD28 antibodies. For human studies, human PBMCs were activated and cultured with Dynabeads™ Human T-Activator CD3/CD28 according to manufacturer’s instruction. T cells were passaged every 48 hr.

### Flow Cytometry Analysis

Single cell suspensions were obtained from cell culture or mouse tissues. Mouse tumors were dissociated by Collagenase (1 mg/mL) and DNAse I (0.2 mg/mL) and lymph nodes were dissociated with 70 μm cell strainer. For each single cell suspension, Fc receptor was blocked with anti-FcγIII/II (clone 2.4G2) for 20 minutes, followed by staining with selective antibodies of cell surface markers and live/dead dyes. Intracellular markers including active caspase 3 and TCF-1 were stained after cell permeabilization with True-Nuclear transcription factor buffer set (BioLegend). Data were collected on BD LSR Fortessa or Beckman CytoFLEX flow cytometer and analyzed by FlowJo (Tree Star Inc., Ashland, OR) software.

### RNA Extraction and Quantitative Real-Time PCR Analysis

Total RNA from T cells was extracted with RNeasy Plus Mini Kit and reversed transcribed with iScript-gDNA Clear cDNA Synthesis Kit (Bio-Rad). Real-time PCR was performed with QuantiFast SYBR Green PCR Kit (Qiagen) according to the manufacturer’s instructions and different primer sets on CFX Connect Real-Time PCR Detection System (Bio-Rad). The levels of gene expression were determined with delta-delta Ct method using β-actin as the internal control.

### Metabolomics analysis

Metabolites were extracted from cultured CD8^+^ T cells with 80/20 methanol/water after 3 cycles of freeze-thaw. Data acquisition was performed by reverse-phase chromatography on a 1290 UHPLC liquid chromatography (LC) system interfaced to a high-resolution mass spectrometry (HRMS) 6550 iFunnel Q-TOF mass spectrometer (MS) (Agilent Technologies, CA). Analytes were separated on an Acquity UPLC® HSS T3 column (1.8 μm, 2.1 × 150 mm, Waters, MA) at room temperature. Mobile phase A composition was 0.1% formic acid in water and mobile phase B composition was 0.1% formic acid in 100% ACN. The LC gradient was 0 min: 1% B; 5 min: 5% B; 15 min: 99% B; 23 min: 99% B; 24 min: 1% B; 25 min: 1% B. The flow rate was 250 μL/min. The sample injection volume was 5 μL. The MS was operated in both positive and negative (ESI+ and ESI-) modes. ESI source conditions were set as follows: dry gas temperature 225 °C and flow 18 L/min, fragmentor voltage 175 V, sheath gas temperature 350 °C and flow 12 L/min, nozzle voltage 500 V, and capillary voltage +3500 V in positive mode and −3500 V in negative. The instrument was set to acquire over the full m/z range of 40–1700 in both modes, with the MS acquisition rate of 1 spectrum/s in profile format. Raw data were processed using Profinder B.08.00 SP3 software (Agilent Technologies, CA) with an in-house database containing retention time and accurate mass information on 600 standards from Mass Spectrometry Metabolite Library (IROA Technologies, MA) which was created under the same analysis conditions. The in-house database matching parameters were: mass tolerance 10 ppm; retention time tolerance 0.5 min. Peak integration result was manually curated in Profinder for improved consistency and exported as a spreadsheet. Data were filtered and analyzed with Metaboanalyst 5.0 and plotted with R 4.0.1. Cut-off for significant change was set to FDR 0.1, fold change 1.5.

### Stable isotope labeling

Activated CD8^+^ T cells from OT-1 mice were rinsed in PBS, then cultured with medium containing the isotopically labeled glucose or lactate for 8 h. For analysis of intracellular metabolites by GC-MS, 2×10 ^6^ cells were rinsed in ice-cold normal saline and lysed with three freeze-thaw cycles in cold 80% methanol. The lysates were centrifuged to remove precipitated protein, and the supernatants with an internal control (10 ul of sodium 2-oxobutyrate) were evaporated, then re-suspended in 40 ul anhydrous pyridine at 70 °C for 15 min. The samples were added to GC/MS autoinjector vials containing 70ul of N-(tert-butyldimethylsilyl)-Nmethyltrifluoroacetamide (MTBSTFA) derivatization reagent and incubated at 70 °C for 1 h. The samples were analyzed using either an Agilent 6890 or 7890 gas chromatograph coupled to an Agilent 5973N or 5975C Mass Selective Detector, respectively. The mass isotopologues were corrected for natural abundance.

### Immunoblot analysis

Total protein from T cells was extracted with SDS lysis buffer (1% SDS, 10 mM HEPES, pH 7.0, 2 mM MgCl_2_, 20 U/mL universal nuclease added protease inhibitor) and quantified with BCA assay (23227, Thermo Fisher). Protein was denatured with Laemmli protein sample buffer with 10% 2-mercaptoethanol by incubating at 95 °C for 10 min. Same amount of denatured protein were subjected to electrophoresis using the standard SDS-PAGE method with 4%-15% gel (4568086, Bio-Rad) and then wet-transferred to a 0.45μm polyvinylidene difluoride membrane. After blocking with 5% non-fat milk, membrane was incubated for 2 h with primary antibodies (1:1,000 dilution with 5% BSA in TBST) at room temperature and for 1 h at room temperature with fluorescent-conjugated secondary antibodies (1:10000 dilution with 5% non-fat milk in TBST). Blots were imaged with Licor Odyssey and analyzed with Image Studio Lite.

### CUT&RUN (Cleavage Under Targets and Release Using Nuclease) PCR

CUT&RUN was performed with CUT&RUN Assay Kit (Cell Signaling Technology #86652) according to the manufacturer’s protocol. Briefly, 1×10^6^ CD8^+^ T cells from OT-1 mice were captured by Concanavalin A-coated magnetic beads (10 μL per million cells) at room temperature for 5 min with rotation. Captured cells were permeabilized in 100 μL Antibody Binding Buffer (+ Spermidine + Protease inhibitor cocktail + digitonin). After permeabilization, 2 μL H3K27ac antibody or 5 μL IgG were added to each reaction for antibody binding (4°C, 6 h, with rotation). After 3 times of washing, cells were incubated with Protein A and Protein G-fused Micrococcal Nuclease (pAG-MNase) at 4°C for 1 h with rotation. By adding cold CaCl_2_, pAG-MNase were activated and antibody-associated protein-DNA complex was released at 4°C for 30 min. Stop solution was then added to the reaction followed by 30 min additional incubation at 37°C. The released DNA was purified using spin columns (Cell Signaling Technology #86652). *Tcf7* super enhancer were quantified with qPCR. The levels of H3K27ac enrichment of *Tcf7* super enhancer were determined with by ratio to IgG sample after normalization to *Cd3e*.

### Quantification and Statistical Analysis

Data were analyzed using GraphPad Prism 8.3. Two-way ANOVA was used to analyze the tumor growth data, logrank test was used to analyze mice survival data, unpaired two-tailed t tests were used to analyze other data unless specified. p < 0.05 was considered statistically significant (*p < 0.05, **p < 0.01, ***p < 0.001 and ****p < 0.0001).

## Acknowledgments

We thank Dr. Ralph DeBerardinis for discussions on metabolomic analysis, Zhaohui Wang, Gang Huang and Zachary Bennett for the helpful discussions on animal experiments. We thank Drs. Jianfeng Ye and Hongyi Zhang for suggestions on sc-RNAseq library preparation and data analysis. Metabolomics analysis was performed by the Metabolomics Facility at the Children’s Research Institute of UT Southwestern Medical Center.

## Author contributions

J.G. and Q.F designed the study. Q.F., Z.L., T.H., J.C., J.Wang, S.L., W.L., Z.S., J.S. performed the experiments. X.Y., Q.F. and J.S. analyzed the multi-omics data. J.G., Y.-X.F., B.L., B.S. supervised the study. Q.F. wrote the original manuscript. J.G., J.Wilhelm, Y.-X.F., B.L., B.S. edited the manuscript. ^†^ These authors contributed equally to this work

## Data and materials availability

All data are available in the main text or the supplementary materials. Processed scRNAseq data with or without lactate treatment can be accessed through the link provided in TableS1 ^‡^ Requests for materials should be sent to Dr. Jinming Gao.

---

**Supplementary Fig. 1.**
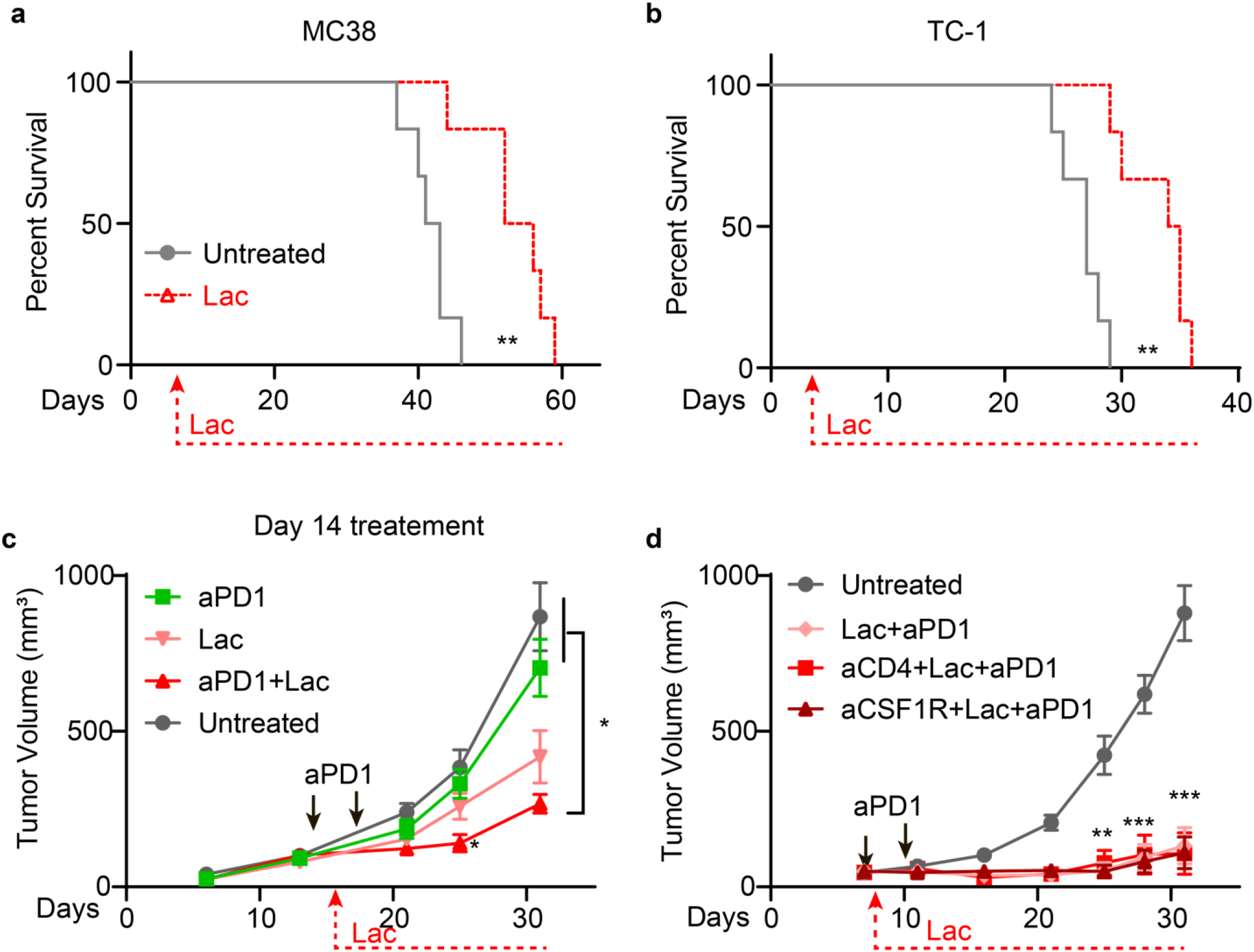
Tumor growth curves of lactate monotherapy and combination therapy with immune cell depletion. Survival curves of lactate treatment in the MC38 (**a**) and TC-1 (**b**) tumor models. C57BL/6 mice (n = 6) were inoculated with 1×10^6^ MC38 tumor cells and treated by lactate. The dose and treatment regimen of lactate is the same as **Fig. 2b**. The untreated group used the same data in **Fig. 2b** for a side-by-side comparison. C57BL/6 mice (n = 6) were inoculated with 1.5×10^5^ TC-1 tumor cells and treated with lactate. The dose and treatment regimen of lactate is the same as **Fig. 2c. c**, Tumor growth curve of MC38 tumor model treated with lactate and anti-PD1 from day 14. C57BL/6 mice (n = 5) were inoculated with 1×10^6^ MC38 tumor cells and treated by lactate. The dose of lactate is the same as **Fig. 2b**. The initial dose of anti-PD1 was on day 14. **d**, CD4^+^ T and macrophage cell depletion assays in the MC38 tumor model. C57BL/6 mice (n = 6) were inoculated with 1×10^6^ MC38 tumor cells and treated with anti-PD-1 (10 mg/kg, day 7 and 10) in combination with glucose or lactate. Anti-CD4 (10 mg/kg) or anti-CSF1R (20 mg/kg) was administered from day 3 and then every three days until the end of the experiment. The dose and treatment regimen of lactate and anti-PD-1 are the same as **Fig. 2e**. The untreated and aPD1+Lac group used the same data in **Fig. 1e** for a side-by-side comparison. Data are shown as means ± SEM. p value was determined by two-way ANOVA, * p<0.05, ** p< 0.01, *** p< 0.001, **** p<0.0001.

**Supplementary Fig. 2.**
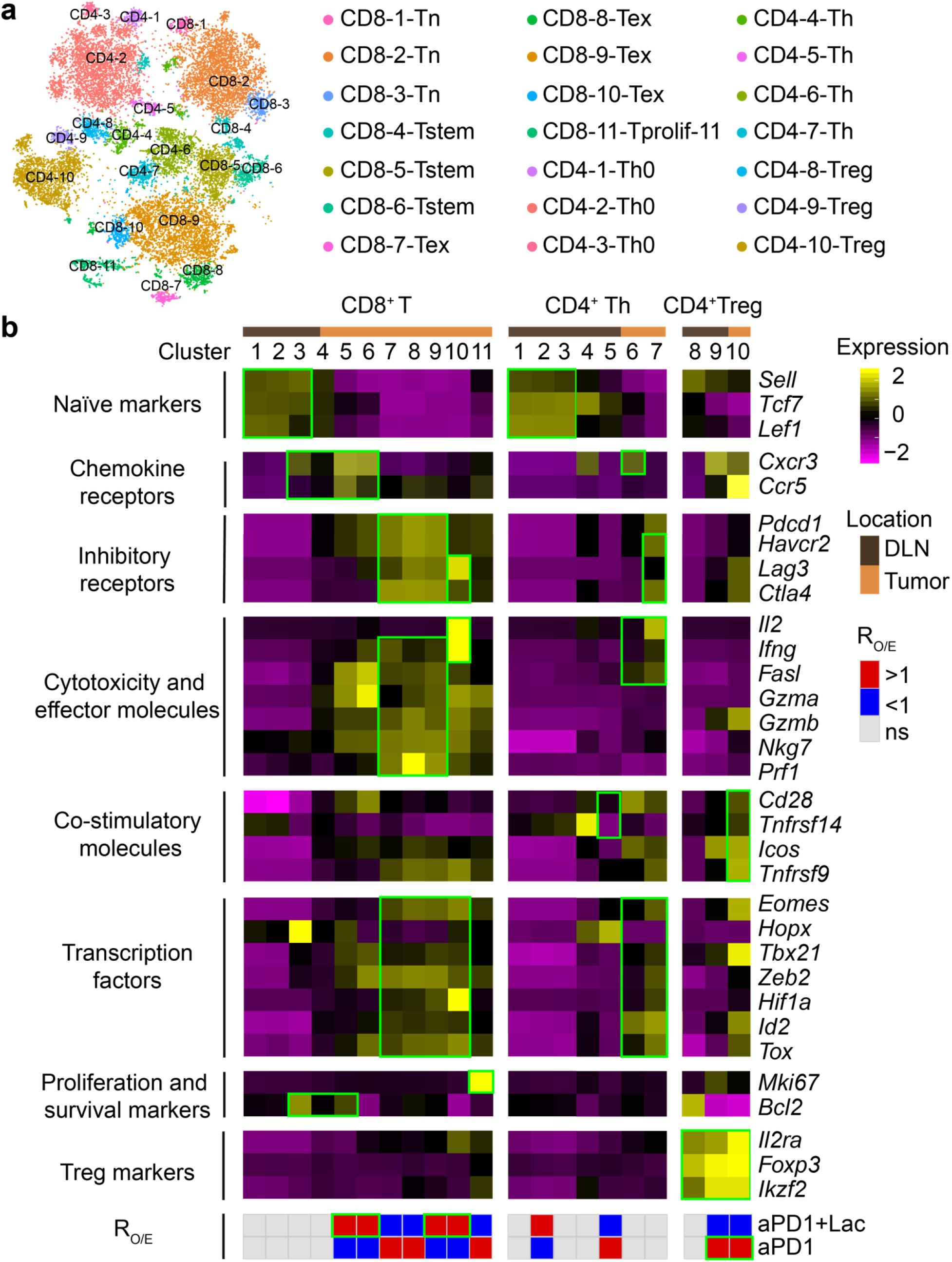
Clustering of T cells from single cell RNA sequencing analysis. **a**, The tSNE projection of 21, 826 single T cells from 2 treatment groups, showing the formation of 21 main clusters, including 11 for CD8^+^ T cells, 7 for conventional CD4^+^ T helper cells (Th: 1-7 of CD4^+^ clusters) and 3 for regulatory T cells (Treg: 8-10 of CD4^+^ clusters). Each dot corresponds to one single cell, colored according to cell cluster. **b**, Mean expression of selected T cell function-associated genes in each cell cluster. Green boxes highlight the prominent patterns defining known T cell subtypes. The ratio of observed cell number to expectation (R_O/E_) of Lac effect on anti-PD-1 treatment for each cluster was calculated by Chi-square test.

**Supplementary Fig. 3.**
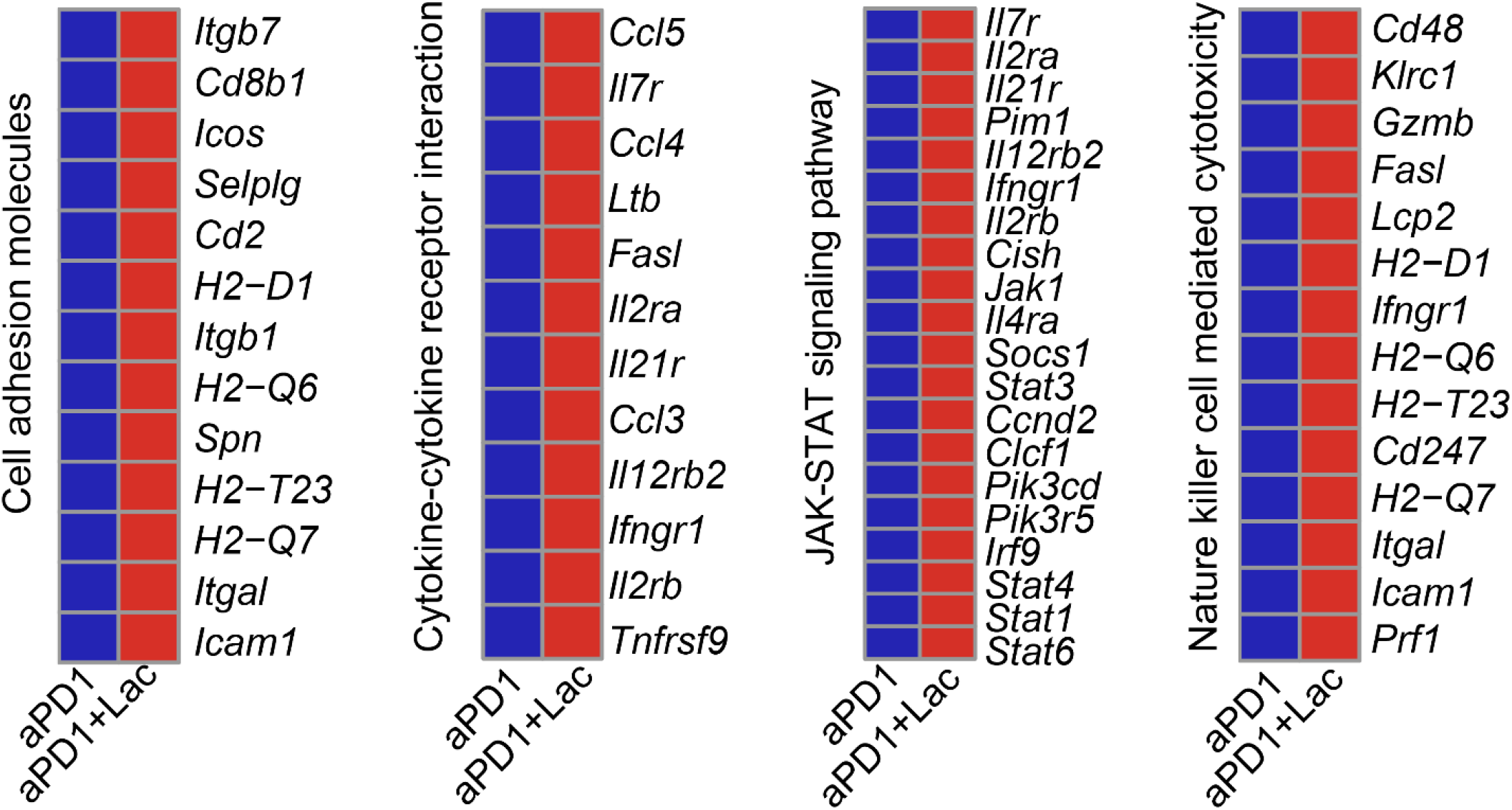
Differentially expressed genes from GSEA analysis in Figure 3.

**Supplementary Fig. 4.**
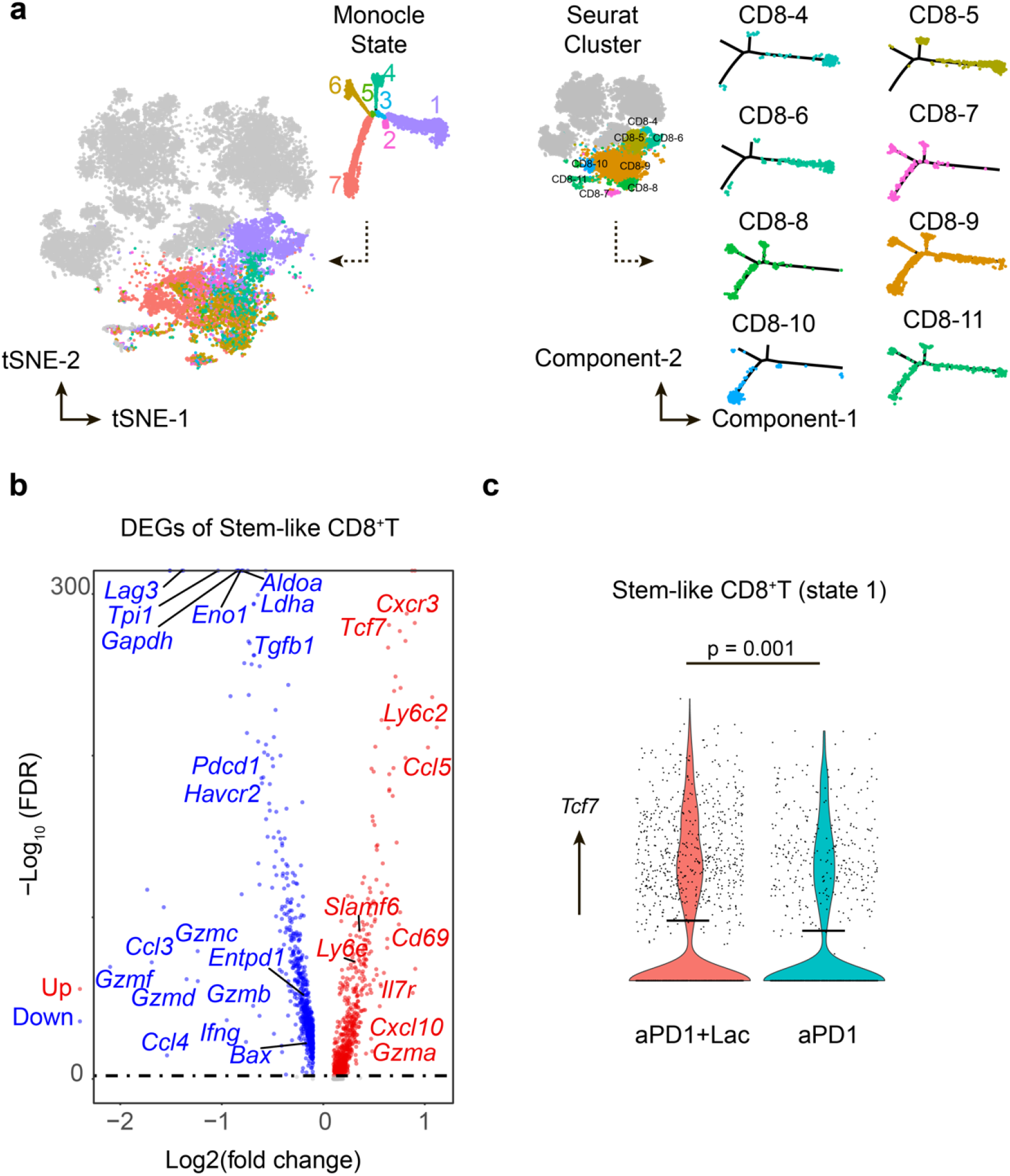
Delineation of stem-like CD8^+^ T cell population by pseudotime analysis. **a**, Correlation of Seurat clusters and monocle states. Stem-like T cells in state 1 are mostly from Seurat cluster CD8-4, CD8-5, CD8-6 and part of CD8-9. **b**, Volcano plot of differentially expressed genes by stem-like CD8^+^ T cells. **c**, *Tcf7* gene expression in stem-like CD8^+^ T cells. p value was determined by Mann Whitney U test.

**Supplementary Fig. 5.**
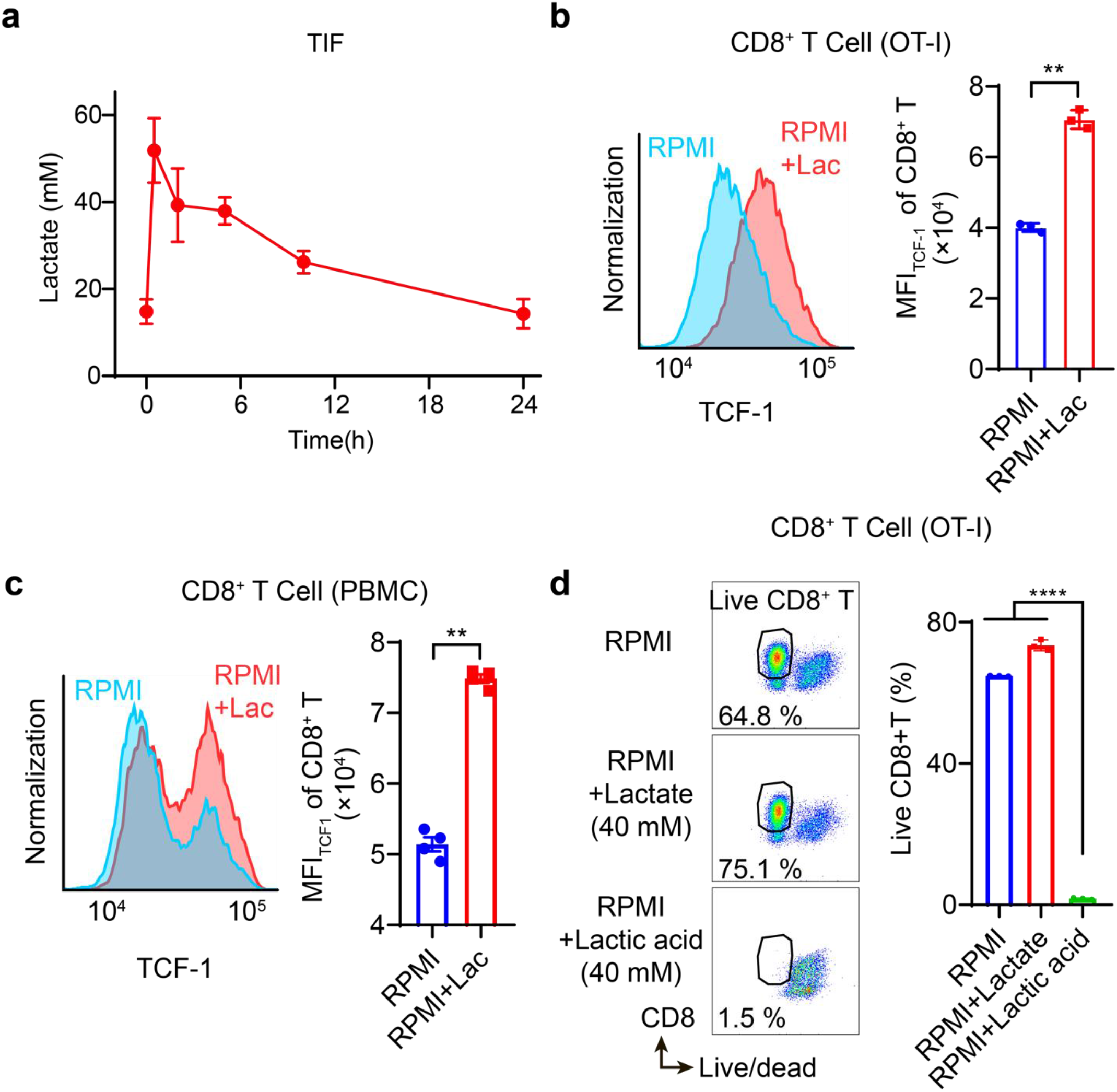
Lactate increases the expression of TCF-1 *in vivo* and *ex vivo* while lactic acid induces T cell death. **a**, Lactate concentration in tumor interstitial fluids (TIF) after subcutaneous lactate injection (n = 3). **b**, Flow cytometry plot and quantification of TCF-1 expressions in *ex vivo* cultured CD8^+^ T cells from OT-I mice on day 4. **c**, Flow cytometry plot and quantification of TCF-1 expressions in *ex vivo* cultured CD8^+^ T cells from human PBMCs on day 4. **d**, Flow cytometry plot and quantification of live CD8^+^ T cells in *ex vivo* cultured splenocytes from OT-I on day 2. Data are shown as means ± SEM. p value was determined by Student’s t test, ** p< 0.01, **** p<0.0001.

**Supplementary Fig. 6.**
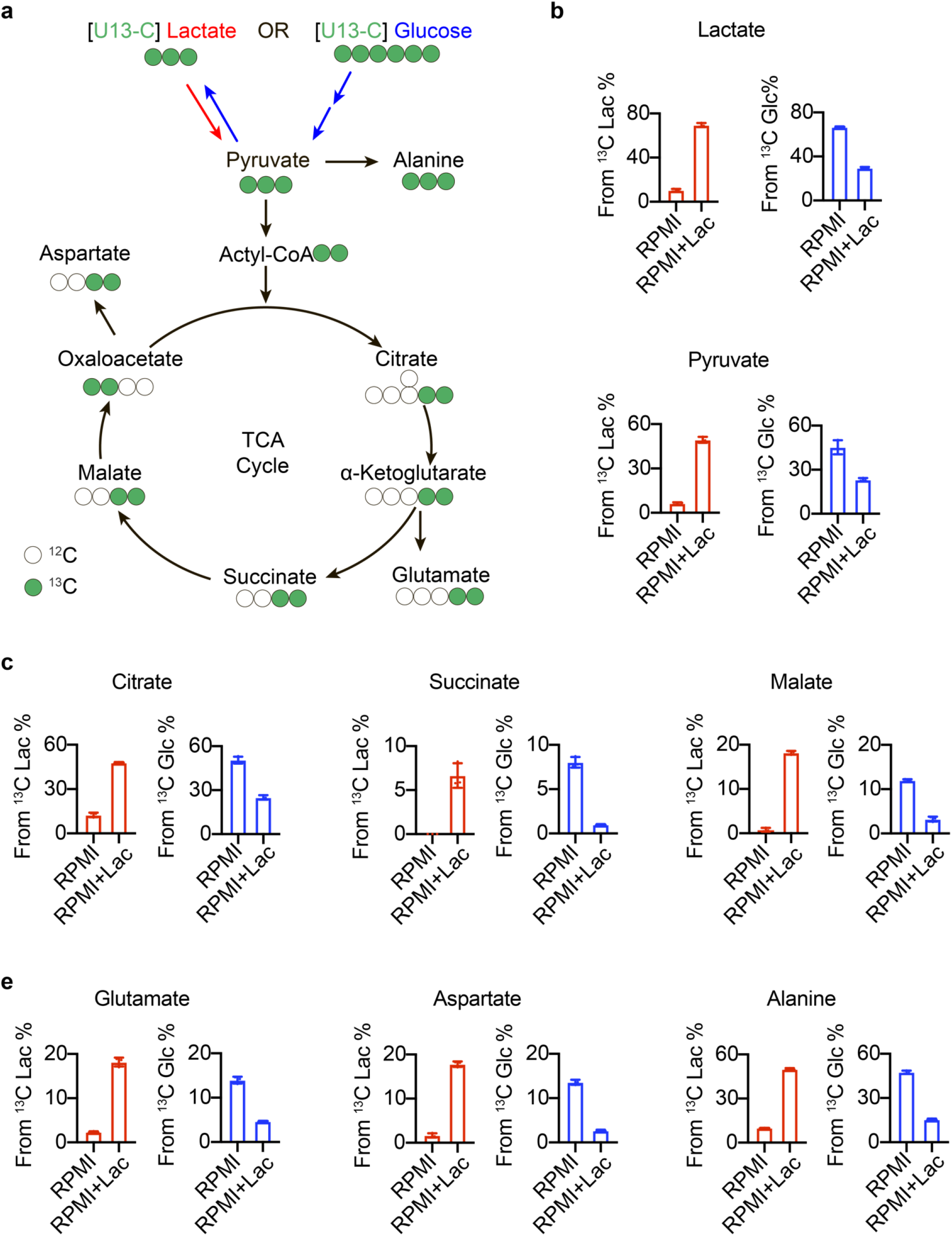
Lactate feeds TCA cycle during *ex vivo* culture of CD8^+^ T cell. **a**, Schematic illustration of isotopologues generated from ^13^C-glucose or ^13^C-lactate. Normalized abundance of intracellular lactate and pyruvate (**b**), TCA cycle intermediates (**c**) and derived amino acids (**d**). Data are shown as means ± SEM. n = 3.

